# Oncogenic c-Myc induces replication stress by increasing cohesins chromatin occupancy

**DOI:** 10.1101/2021.07.25.453647

**Authors:** Silvia Peripolli, Tanya Singh, Harshil Patel, Leticia Meneguello, Koshiro Kiso, Peter Thorpe, Cosetta Bertoli, Robertus A.M. de Bruin

## Abstract

Oncogene-induced replication stress is a major driver of genomic instability in cancer cells, with a central role in both cancer initiation and progression (1). Despite its critical role in cancer development, the mechanisms that lay at the basis of oncogene-induced replication stress remains poorly understood. Here, we investigate the mechanism of c-Myc-induced replication stress. Our data shows that c-Myc induces replication stress by increasing the amount of cohesins bound to chromatin in the G1 phase of the cell cycle. This is independent of previously suggested mechanisms involving deregulation of replication initiation and transcriptional interference. Restoring the amount of chromatin-bound cohesins to control levels, or preventing the accumulation of cohesins at CTCF sites, in cells experiencing oncogenic c-Myc activity prevents replication stress. Increased cohesins chromatin occupancy correlates with a c-Myc-dependent increase in the levels of the cohesion loader Mau2. Preventing c-Myc-induced increase in Mau2 reduces oncogene-induced replication stress. Together our data support a novel mechanism for oncogene-induced replication stress. Since c-Myc activation is a crucial event in many human cancers (2), identifying the mechanisms through which this oncogene promotes replication stress provides critical insights into cancer biology.

---

Replication stress (RS) is defined as the slowing-down and/or stalling of DNA replication forks and is a major source of genomic instability in cancer cells, being often caused by activation of oncogenes or loss of tumour suppressors (3), (1, 4, 5). Despite the critical role of oncogene-induced RS in cancer, the mechanisms that generate it remain unclear (Figure 1a). Different oncogenes have been studied in various systems, and the general consensus is that RS is likely the result of deregulation of replication initiation events (6). Other proposed mechanisms involve the interference between the replication and transcription machineries (7–9). In the case of oncogenic overexpression of Cyclin E, different mechanisms have been reported (7, 10, 11). The reduced length of G1 following Cyclin E overexpression has been associated with a decrease in licensing events, which results in fewer replication complexes available for replication. This is thought to culminate in genomic instability due to under-replication (12). More recently, Cyclin E overexpression has also been shown to cause RS by increased transcription-replication collisions in transcribed genes (7, 11). In contrast, the oncogenic activity of the transcription factor c-Myc has been reported to increase replication initiation events, thus causing over-replication (4, 5). Surprisingly, while c-Myc is thought to induce a large transcriptional program to promote proliferation and growth (13), it has not been linked to increased transcription-replication interference.

**Fig. 1.**
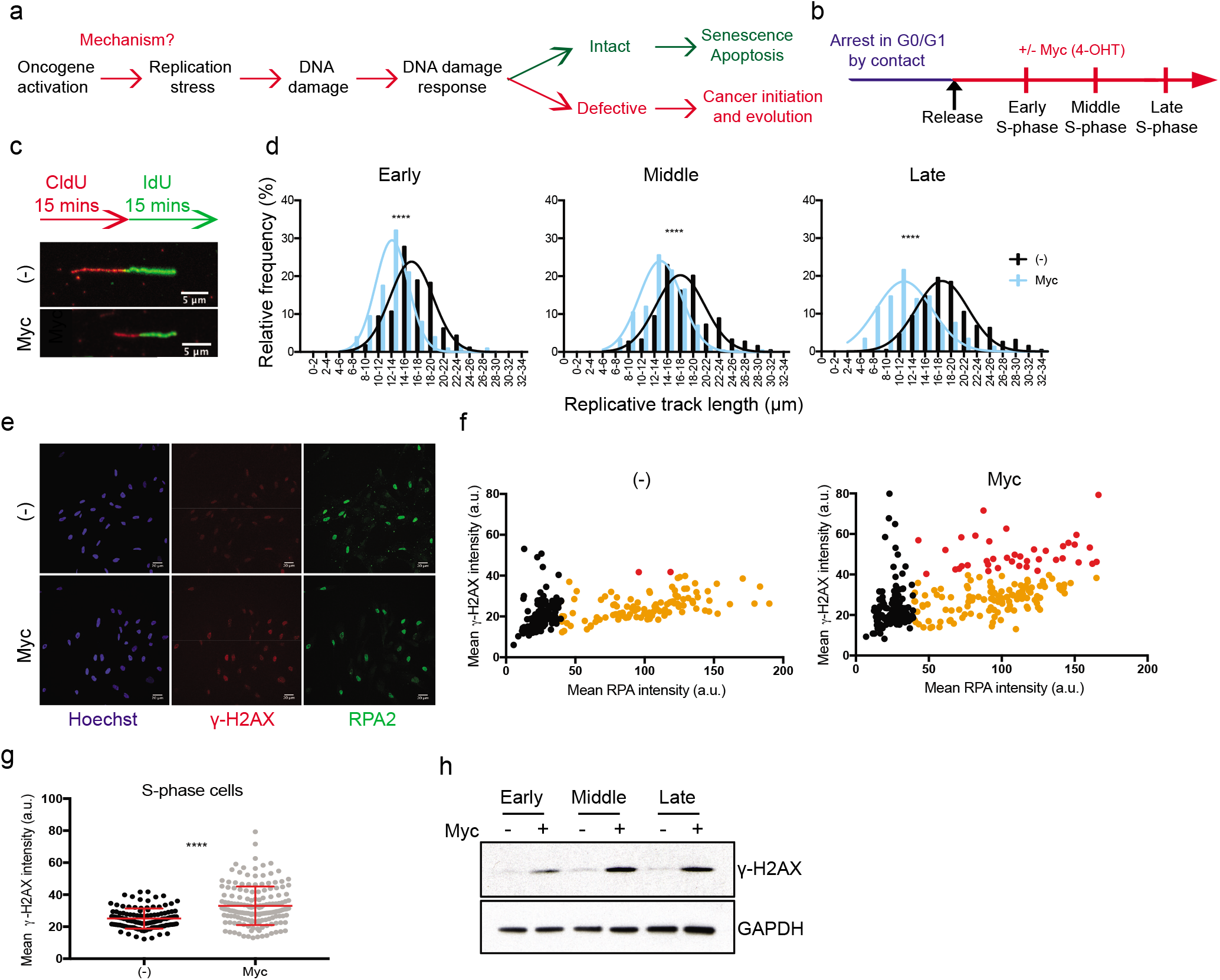
c-Myc-induced replication stress and DNA damage depend on events in G1 phase. a) Schematic of known oncogene-induced replication stress mechanisms. b) Schematic of the synchronisation experiments for G1-S release. RPE1 c-Myc ER cells were left to grow to confluence, then trypsinised and plated in fresh medium. After cell spreading, 4OH-T was added to induce c-Myc or left untreated as control. c) DNA fibre analysis of synchronised cells. Schematic showing the pulse labeling with the two nucleotide analogues. Immunofluorescence of representative fibres. d) Histograms reporting the distribution of fibre length for control and c-Myc-induced cells at the reported times after release from arrest; early=18 hr, middle=22 hr, late=24 hr; p-value****<0.0001 calculated with Mann-Whitney test. Representative of n=2 experiments. e) Representative images of RPA and *γ*H2AX immunofluorescence. f) Immunofluorescence staining of chromatin-bound RPA and *γ*H2AX after release from confluence arrest. Scatter plot showing the intensity of RPA and *γ*H2AX signal in single pre-extracted nuclei. Black=RPA negative cells, orange=RPA positive cells, red=RPA positive cells with higher *γ*H2AX signal. Representative of n=2 experiments. g) Graph showing *γ*H2AX intensity in individual S phase cells plotted in f). p-value****<0.0001 calculated with Mann Whitney test. Representative of n=2 experiments. h) Western blot of *γ*H2AX at the indicated time-points after release from G1 arrest, with and without c-Myc activation; early=18 hr, middle=22 hr, late=24 hr; GAPDH is a loading control. Representative of n=3 experiments.

Besides transcription machineries other large protein complexes bound to DNA could interfere with replisome progression. The cohesin complex is probably one of the most abundant protein complexes interacting with the DNA. Cohesins are ring-shaped multiprotein complexes comprising two major subunits, Structural Maintenance of Chromosome (Smc) 1 and Smc3, along with the kleisin subunit Rad21 and Stag1 and Stag2 in mammalian cells (14). The loading of cohesins onto DNA is highly regulated, and in mammalian cells depends on the activity of the loaders Mau2 and Nipped-B-like protein (NIPBL). The activity of the loaders is antagonised by the release factor Wapl (15). While loading and release occur throughout the entire cell cycle, during S-phase and G2 cohesins interaction with the DNA is more stable (16). This is due to the establishment factors, Esco1 and Esco2 in mammalian cells, which acetylate the Smc3 subunit of the cohesin complex (17), and thus prevent the release activity of Wapl (18).

Recent evidence confirmed that in mammalian cells, as previously reported in budding yeast (19), cohesin rings are able to move along the DNA in a transcription-dependent manner (20). Binding of the CCCTC-Binding Factor (CTCF) to CTCF sites is involved in the organisation of spatially interacting regions of chromatin (21), known as topologically associated domains (TADs) (21, 22). It has been shown that CTCF sites can act as a road-block for cohesins, with accumulation of cohesins often detected at CTCF sites in mammalian cells (23, 24). While the majority of cohesins interact dynamically with the DNA, some are associated more stably, and reside at CTCF sites, where they participate in the organization of chromatin loops (25).

To establish which of the different mechanisms contribute to c-Myc-induced RS, we exploited a c-Myc-ER inducible RPE1 hTERT cell line (26). In this system oncogenic c-Myc activation depends on the translocation of the c-Myc-ER protein into the nucleus after 4OH tamoxifen (4OH-T) addition (Figure S1a, S1b), which is independent of c-Myc-ER protein levels (Figure S1c). 24 hr to 48 hr after 4OH-T addition oncogenic c-Myc activity can be observed via a decrease in colony formation (S1d) and increased gene expression of well-known c-Myc target genes at the mRNA (Figure S1e, S1f) and protein level (Figure S1g). As shown previously (26), activation of c-Myc-ER by 4OH-T induces RS-induced DNA damage response within 24 hours (Figure S1i, S1j) and shortening of DNA fibre length (Figure S1k, S1l), indicative of slowing down or replication forks, supporting the use of this system to study c-Myc-induced RS.

To gain some initial insights into how c-Myc induces RS, we arrested cells in different cell cycle phases, released them with or without c-Myc activation and tested the levels of RS and DNA damage in the following S phase. First, we used confluency to arrest cells in G0/G1. After release into the cell cycle, the oncogene c-Myc was activated by adding 4OH-T or left untreated as control (Figure 1b, S1m), allowing us to study the first G1 and S phases after the oncogene activation. In order to measure RS, we analysed the length of DNA fibres. The activation of c-Myc reduces the average DNA fibre length compared to control, suggestive of slowing down of replication forks (Figure 1c, 1d and Figure S1n). We then measured DNA damage by monitoring the phosphorylation of the histone H2AX. We observed increased levels of the DNA damage marker *γ*H2AX in S phase cells with both Western blot and immunofluorescence (Figure 1e-h). We confirmed these findings by synchronising the cells via nocodazole shake-off (Figure S1o-s). These data suggest that the activation of c-Myc during G1 and S phase can induce RS and DNA damage.

To test if G1 phase is required for c-Myc to induce RS, we synchronised cells in early S phase by adding hydroxyurea (HU). Subsequently we washed out HU to allow S phase progression with or without c-Myc activation and analysed the levels of RS and DNA damage (Figure S2a, S2b). Surprisingly, we did not observe any decrease in DNA fibre length in c-Myc activated cells compared to control (Figure S2c). On the contrary DNA fibres appear longer upon c-Myc induction, indicating that c-Myc activity during S phase does not cause RS, but might even increase the replication capacity of the cell as reported in Pennycook et al (27). Correspondingly, we did not observe any increase in DNA damage in c-Myc cells (Figure S2d). As expected, c-Myc activation increased replication initiation events (origin firing, Figure S2c) and increased Cyclin E levels, a transcriptional target of c-Myc, as well as Mau2 levels (Figure S2e, S2f). This data suggests that c-Myc-induced increase in origin firing does not cause RS per se. Since HU treatment by itself causes RS, which might mask c-Myc-induced RS, we also released cells synchronously into S phase after a G1 arrest via CDK4/6 inhibition by Palbociclib, as reported by Trotter et al (28). c-Myc was activated, via addition of 4OH-T, either immediately after release, therefore throughout G1 phase, or immediately before entering S phase, and DNA damage and DNA fibre length were analysed as above (Figure S2g-j). These data confirm that c-Myc activity during G1 is required to generate RS in S phase.

A possible cause could be a decrease in G1 length, which has been associated with reduced origin licensing and consequent RS upon Cyclin E overexpression (12). We evaluated replication origin licensing in pre-extracted samples by measuring chromatin bound Mcm7, a component of the helicase complex that is loaded on the DNA during G1 as in Ekholm-Reed et al (12). We did not observe a decrease in origin licensing, (Figure S2k), indicating that reduced origin licensing is an unlikely causative mechanism for c-Myc-induced RS.

It has previously been reported that RS can result from increased transcription-replication collisions (7, 29). To test if interference between replication and c-Myc-induced transcription contributes to RS we decreased global transcription levels after c-Myc activation via treatment with the RNA polymerase inhibitor 5,6-Dichloro-1-β-d-Ribofuranosyl Benzimidazole (DRB) for 2 hr, like in Kotsantis et al (8), during the first S phase after release from G1 (Figure S2l-m). We confirmed that c-Myc activation increases global transcription levels, by 5-Ethynyl Uridine (EU) incorporation, and that DRB reduces it, (Figure S2l). However, DNA fibre analysis shows that this does not rescue the c-Myc-dependent decrease in DNA fibre length (Figure S2m). This suggests that in this system transcription-replication interference does not significantly contributes to c-Myc-induced RS levels, though we cannot exclude a role for this mechanism in other settings. We next focused on alternative mechanisms that could contribute to c-Myc-induced RS. A potential source of RS could be protein complexes interacting with the DNA during S phase that could slow down fork progression. A potential candidate for this would be the cohesin complexes. Whilst cohesin has been shown to have a role in preventing RS-induced DNA damage (30), we hypothesised that a c-Myc-induced increase in cohesin occupancy during G1 could slow down replisome progression during S phase as suggested in Kanke et al (31) and Morales et al (32). We therefore analysed the fraction of chromatin bound cohesin subunits Smc1 and Smc3 by Immunofluorescence (IF) of pre-extracted samples, in cells released from G0/G1 with and without c-Myc. These data show that higher levels of Smc1 and Smc3 are detected on chromatin in c-Myc-activated cells both in G1 and in S phase (Figure 2a, S3a). This was confirmed in asynchronous cell populations (Figure S3b) and by chromatin preparations followed by Western blot analysis (Figure 2b).

**Fig. 2.**
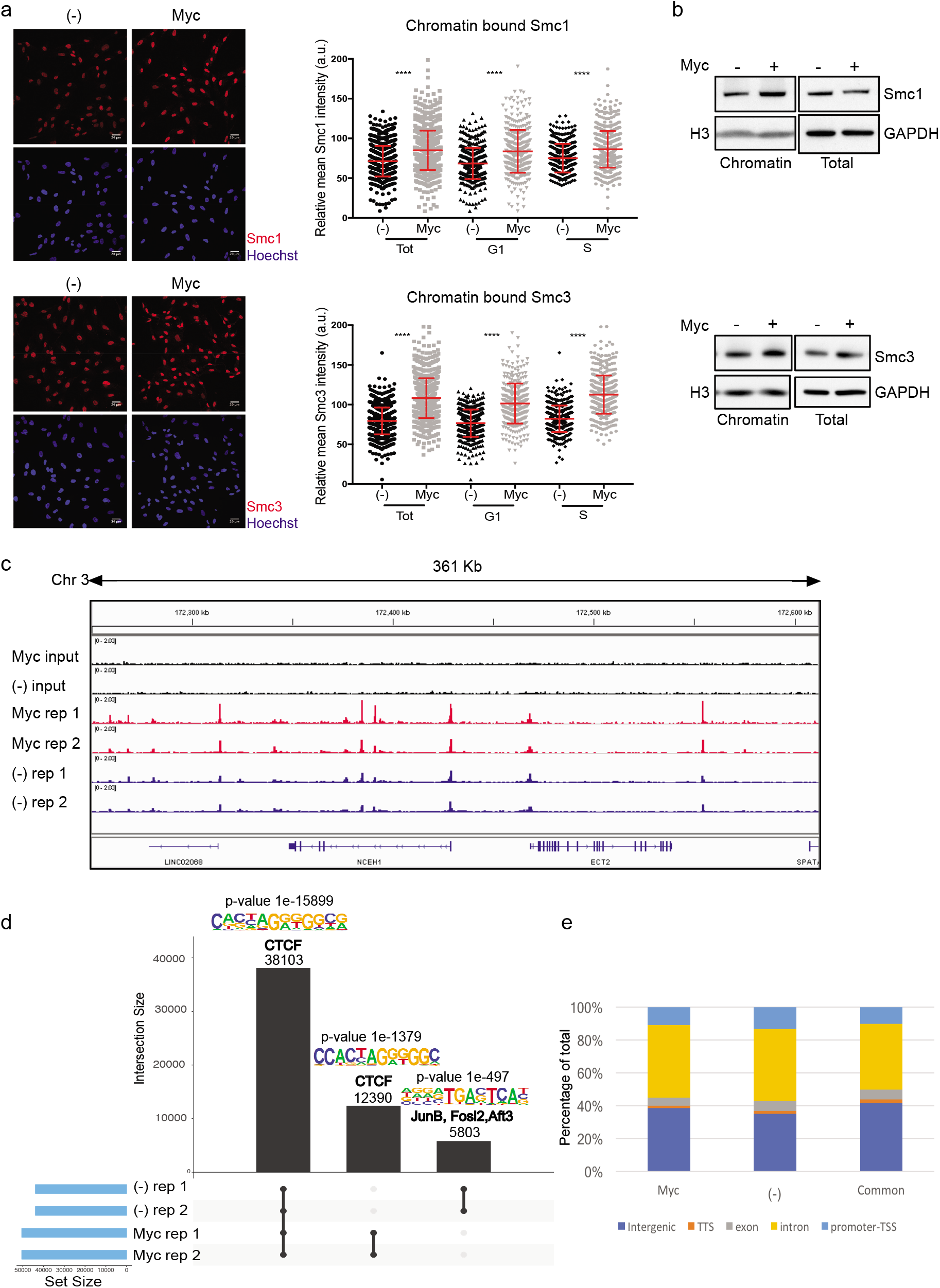
c-Myc activation increases cohesion chromatin occupancy in G1. a) Synchronised cells were released into the cell cycle and immunofluorescence of chromatinbound cohesin subunits Smc1 and Smc3 were performed at 18 hr after release. Left panels; representative images of Smc1, Smc3 and Hoechst. Right: graph reporting the intensity of Smc1 and Smc3 signals in total, S phase and G1 phase single pre-extracted nuclei of untreated and c-Myc cells. p-value****<0.0001 calculated with Mann-Whitney test. Pool of n=3 experiments. b) Western blot of chromatin preparations and total cell lysates in asynchronous population with and without c-Myc activation for 16 hr. GAPDH and H3 are loading controls. Representative of n=3 experiments. c) Binding of Smc1 to the reported DNA loci in untreated and c-Myc-activated cells. Two repeats for each condition are represented. d) Graph representing the analysis of Smc1 binding distribution in c-Myc and untreated cells. Binding motif prediction with p-value for each group, along with the published similar consensus identified. e) Analysis of the genomic features of peak locations in c-Myc only, untreated only and common sites.

Our IF analysis on pre-extracted nuclei together with the chromatin preparation and Western blot analysis establish an overall increase in chromatin occupancy of the cohesin subunits Smc1 and Smc3, but do not reveal where these accumulate in the genome. To investigate if this c-Myc-dependent increase in chromatin bound cohesin accumulates at specific sites, we established genome-wide binding of the cohesin component Smc1 by ChIPseq in cells with activated c-Myc or control (Figure 2c-e). We performed these experiments at 48 hr after c-Myc activation, when we observe consistent levels of RS (Figure S1k), and no obvious differences in cell cycle distribution between control and c-Myc-activated cells (Figure S1h). These experiments show that cohesin accumulates at 50,493 sites in c-Myc cells compared to 43,906 in control (Figure 2d, see set size). In particular, while the vast majority of peaks (38,103) are common to control and c-Myc-activated cells, 12,390 are unique for c-Myc and 5,803 are only present in the control (Figure 2d). Overall, the distribution of cohesin in genic and intergenic regions does not change upon c-Myc activation (Figure 2e). Motif analysis of the peaks in control shows enrichment of CTCF sites, which is in line with published data (23). Enrichment of CTCF sites can be detected in the peaks in common and in the c-Myc-specific peaks. These data suggest that the c-Myc-dependent increase in chromatin bound cohesin primarily accumulates at CTCF sites, in agreement with the reported location of cohesins on DNA (23, 24) (Figure 2d).

Next, we wanted to investigate if the increase in cohesin chromatin occupancy contributes to c-Myc-induced RS. We tested this by reducing the levels of cohesins on DNA in cells experiencing oncogenic c-Myc activity by targeting the cohesin component Rad21. We used a non-efficient siRNA to ensure that the reduction of Rad21 would not cause cell cycle defects during the first S phase (Figure S3c). Analysis of Smc1 binding to chromatin confirms that Rad21 knockdown reduces the levels of chromatin-bound cohesins both in untreated and in c-Myc cells (Figure 3a and b). Importantly, the levels of Smc1 on chromatin in Rad21-depleted c-Myc cells are similar to the untreated control, allowing us to test if the increased chromatin binding of cohesins is at the basis of generating RS.

**Fig. 3.**
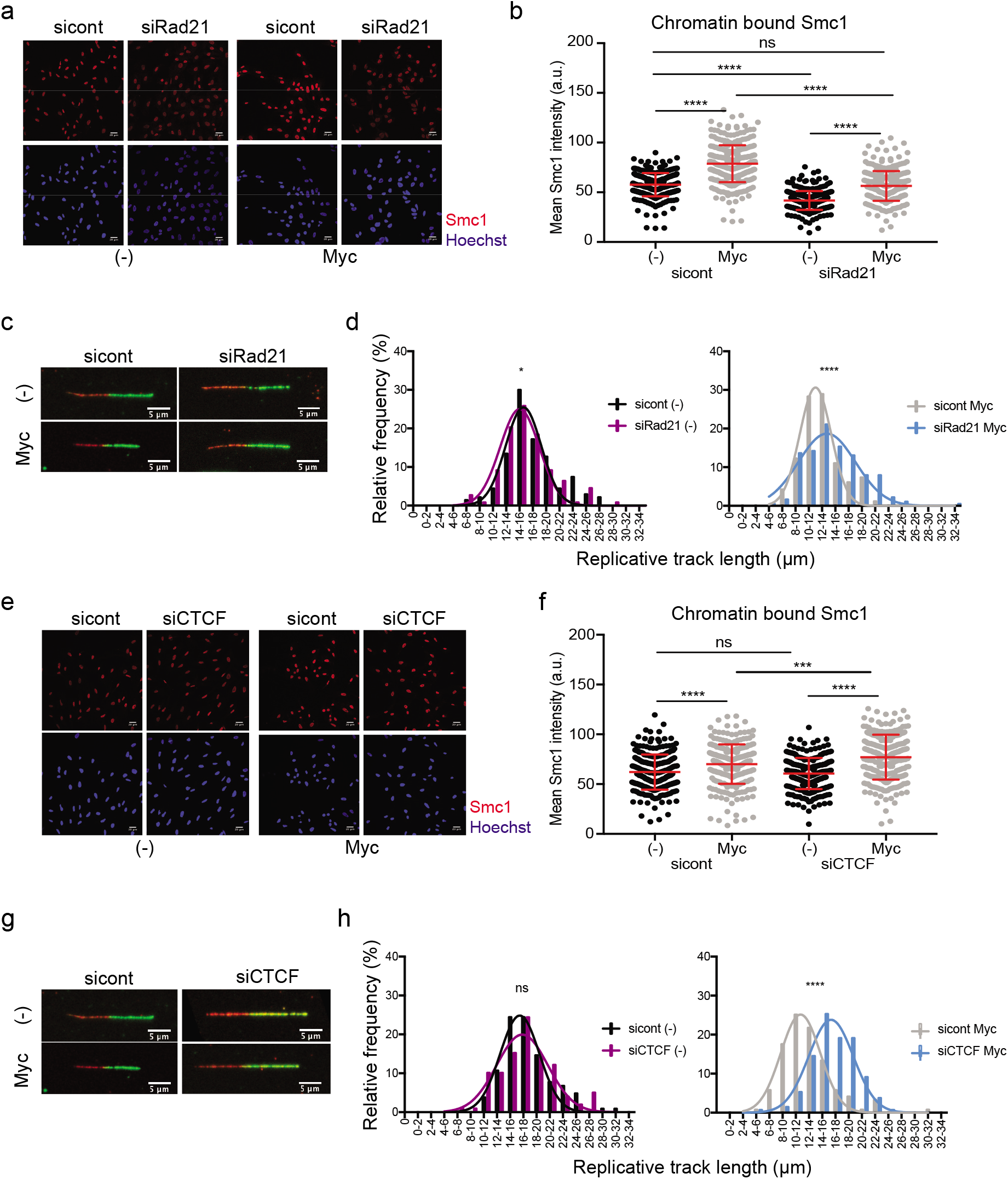
Reducing the levels of cohesin on chromatin at CTCF sites prevents c-Myc-induced replication stress. a) Immunofluorescence showing chromatin-bound Smc1 in synchronised sicontrol and siRad21 depleted cells at 20 hr after release, with and without c-Myc activation. b) Graphs representing the intensity of chromatin-bound Smc1 signal in immunofluorescence. p-value****<0.0001 calculated with Mann-Whitney test. Representative of n=3 experiments. c) Immunofluorescence of representative fibres in synchronised sicontrol and siRad21 depleted cells at 20 hr after release from G1. d) Histograms reporting the distribution of fibre length in synchronised sicontrol and siRad21 depleted cells. p-value****<0.0001 calculated with Mann-Whitney test. Representative of n=3 experiments. e) Immunofluorescence showing chromatin-bound Smc1 in synchronised sicontrol and siCTCF depleted cells at 20 hr after release, with and without c-Myc activation. f) Graphs representing the intensity of Smc1 signal in the immunofluorescence. p-value****<0.0001 calculated with Mann-Whitney test. Representative of n=3 experiments. g) Immunofluorescence of representative fibres in synchronised sicontrol and siCTCF depleted cells at 20 hr after release from G1. h) Histograms reporting the distribution of fibre length in synchronised sicontrol and siCTCF depleted cells. p-value****<0.0001 calculated with Mann-Whitney test. Representative of n=3 experiments.

Reducing the levels of cohesin chromatin occupancy in cells experiencing oncogenic c-Myc increases DNA fibre length in the first S-phase in both synchronous and asynchronous cell populations (Figure 3c and 3d, Figure S3d and S3e). Rad21 depletion did not affect the extent of replication origin firing and the cell cycle profiles (Figure S3f, S3g), nor the expression level of well-established c-Myc target genes (Figure S3h) supporting the idea that it is the increase of cohesins on DNA that contributes to RS. Next, we tested if the c-Myc-dependent accumulation of cohesin specifically at CTCF sites was important for c-Myc-induced RS. Depleting CTCF (Figure S3i), completely rescued RS induced by c-Myc, without reducing the global amount of cohesins on DNA or affecting the cell cycle profiles (Figure 3e-h, S3j, S3k). We finally analysed the RS and DNA damage response activation in these cells, following Rad21 and CTCF depletion. We observed a decrease in both Chk1 phosphorylation and *γ*H2AX levels compared to control silencing upon c-Myc activation, suggesting that in both cases RS-induced DNA damage was reduced (Figure S3l, S3m).

Together these data indicate that the c-Myc-dependent increase in cohesins chromatin occupancy, likely at CTCF sites, causes a slowdown of replication forks to generate RS. To investigate how c-Myc could increase cohesin chromatin occupancy, we tested if its activation affects the expression levels of cohesin subunits and regulators. Smc1, Smc3 and Rad21 protein levels did not change significantly upon c-Myc activation (Figure S4a). Interestingly, protein levels of the cohesin loader Mau2 increased upon c-Myc activation in both synchronised (Figure 4a) and asynchronous cells (Figure 4b, S4b), which corresponds to an increase in mRNA levels (Figure S4c, S4d). In vertebrates, cohesin loading requires NIPBL and Mau2 (33, 34). While NIPBL is the proper cohesin loader, Mau2 stabilises the protein levels of NIPBL (23, 35, 36), therefore we analysed NIPBL levels in c-Myc-activated cells. Both Mau2 and NIPBL protein levels increase upon c-Myc activation (Figure S4b), while NIPBL mRNA does not change significantly (Figure S4d), suggesting that c-Myc increases Mau2 expression that in turn stabilises NIPBL.

**Fig. 4.**
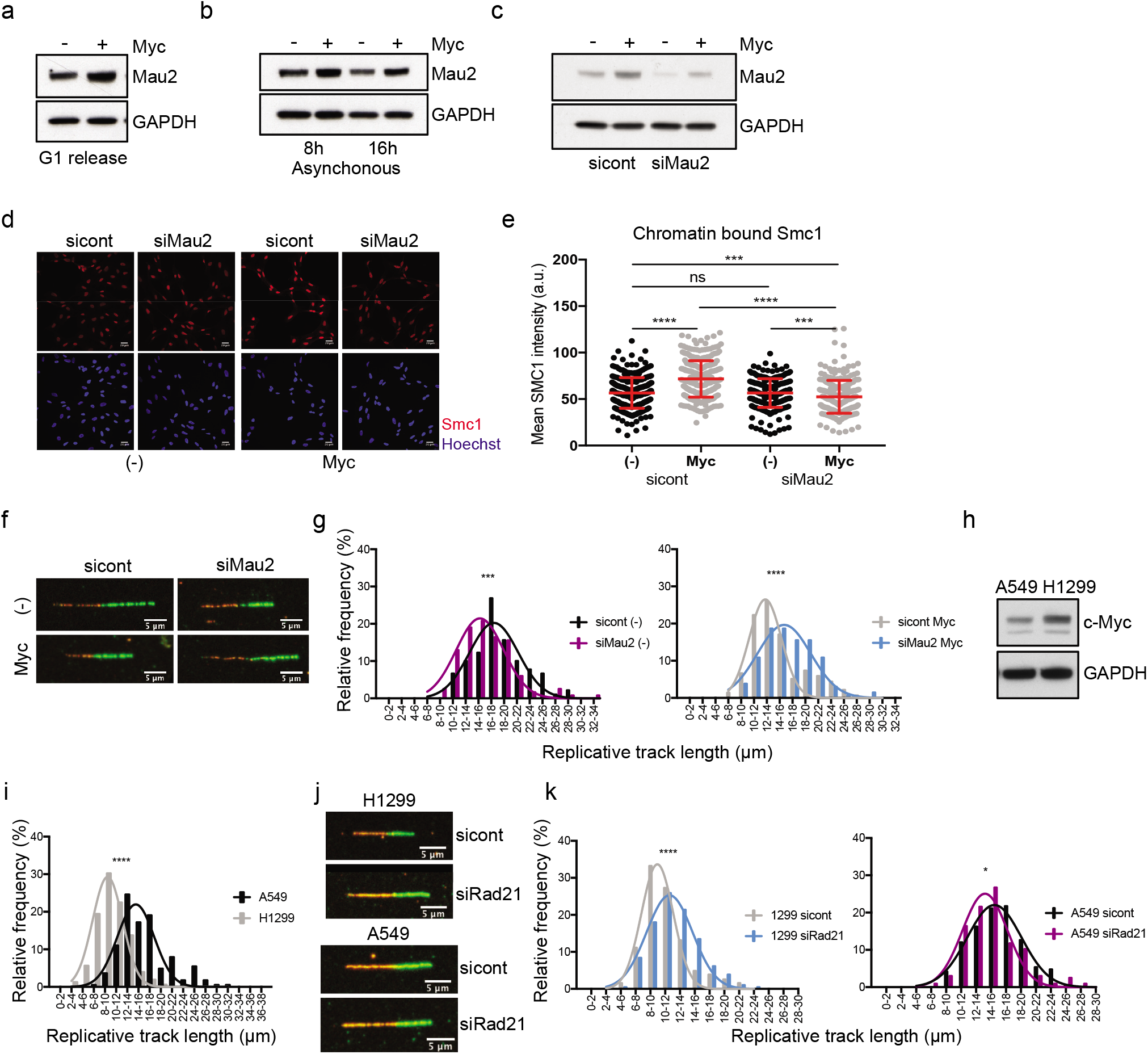
Reducing the levels of cohesin loader Mau2 prevents c-Myc-induced replication stress. a) Western blot showing total levels of the cohesin loader Mau2 in synchronised cells at 20 hr after release, with and without c-Myc activation. Representative of n=3 experiments. GAPDH is a loading control. b) Western blot showing total levels of the cohesin loader Mau2 in asynchronous population with and without c-Myc activation for the indicated timepoints. Representative of n=2 experiments. GAPDH is a loading control. c) Western blot showing Mau2 knock-down in synchronised cells at 20 hr after release, with and without c-Myc activation. Representative of n=3 experiments. GAPDH is a loading control. d) Immunofluorescence showing chromatin-bound Smc1 in synchronised sicontrol and siMau2 depleted cells at 20 hr after release, with and without c-Myc activation. e) Graphs representing the intensity of chromatin-bound Smc1 signal in the immunofluorescence. p-value****<0.0001, ***=0.0001 calculated with Mann-Whitney test. Representative of n=2 experiments. f) Immunofluorescence of representative fibres in synchronised sicontrol and siMau2 depleted cells at 20 hr after release from G1. g) Histograms reporting the distribution of fibre length in synchronised sicontrol and siMau2 depleted cells. p-value****<0.0001 calculated with Mann-Whitney test. Representative of n=3 experiments. h) Western blot showing total levels of c-Myc in A549 and H1299 cells. Representative of n=3 experiments. GAPDH is a loading control. i) Histograms reporting the distribution of fibre length in A549 and H1299 cells. p-value****<0.0001 calculated with Mann-Whitney test. Representative of n=3 experiments. j) Immunofluorescence of representative fibres in A549 and H1299 cells after Rad21 depletion. k) Histograms reporting the distribution of fibre length in A549 and H1299 cells after Rad21 depletion. p-value*=0.0356, ****<0.0001 calculated with Mann-Whitney test. Representative of n=3 experiments.

To verify whether c-Myc-induced increase in cohesin chromatin occupancy could be partially due to increased loading via upregulating Mau2, we reduced Mau2 levels in c-Myc activated cells to control levels using a non-efficient siRNA (Figure 4c) and tested the presence of cohesins on chromatin and RS. Similar to Rad21 reduction, preventing the increase in Mau2 prevented excess cohesin loading onto chromatin (Figure 4d, e) and RS upon c-Myc activation in both synchronised (Figure 4f, g and S4e) and asynchronous cells (Figure S4f and S3l, m). As for Rad21 knock down experiments, the extent of origin firing and cell cycle distribution is not affected by Mau2 depletion (Figure S4g). These data indicate that c-Myc-dependent increase in Mau2 levels could be at the basis of the increased loading of cohesins on chromatin, which causes RS during S phase.

To investigate if the c-Myc-dependent increase in Mau2 levels could contribute to the generation of RS we transiently expressed Mau2-GFP, or GFP as control, in RPE1 cells and analysed RPA phosphorylation, marker of RS, and *γ*H2AX, marker of DNA damage, by quantitative immunofluorescence and western blot (Figure S4h-m). Whilst both markers increase in IF in transfected cells, an increase in phosphorylation of RPA is particularly pronounced in IF and in total lysates. These data suggest that overexpression of Mau2 alone can cause some RS and RS-induced DNA damage.

We finally tested whether this mechanism of c-Myc-induced RS contributes to RS in cancer cells (Figure 4h-k). We selected a couple of lung cancer cell lines expressing different levels of c-Myc and measured the length of DNA fibres to evaluate the presence of RS. We observed reduced fibre length in the H1299 cell line compare to the A549 cells, which correlates with higher levels of c-Myc in H1299 cells (Figure 4h, 4i). To establish if a reduction in cohesins can rescue RS levels we depleted Rad21 (Figure S4n) and measured DNA fibre length in both cell lines (Figure 4j, k and S4o). Rad21 depletion increased fibre length only in the H1299 cells, where c-Myc is highly expressed, but not in A549 cells, supporting a causative role for cohesins in RS, which is in the agreement with our findings. Interestingly depletion of Rad21 in the A549 cells, which do not experience RS, partially reduces fibre length. This suggests a protective role for cohesins in preventing RS, which is in agreement with reported work, supporting a double-edge involvement of cohesins in RS.

Our data show that a c-Myc-induced increase in cohesins on the DNA contributes to the induction of RS. This is different from previously reported mechanisms of oncogene-induced RS, which are linked to deregulation of replication origin usage and/or transcription/replication interference. We show that the c-Myc-dependent accumulation of cohesins on chromatin, most likely at CTCF sites, can cause RS. Our data indicates that the levels of the cohesin loader Mau2 are upregulated by c-Myc activity, and that overexpression of Mau2 alone can cause RS, providing a potential mechanism through which c-Myc affects cohesin regulation. Overall, our data show that excessive cohesins on chromatin can interfere with the progression of replication forks, thus contributing to oncogene-induced RS. Our findings are surprising in light of previous work which indicates an important role for cohesins in preventing RS and DNA damage (37, 38). Based on our data we speculate that whilst the presence of cohesins during S phase is required to protect stalled forks and repair damaged DNA, excessive cohesin accumulation in an oncogenic context can interfere with the progression of the replisome. This is in agreement with recently published work in mammalian cells which shows that increased presence of cohesins on DNA slows down fork progression (32) and with work in yeast, which shows that DNA damage accumulates in SMC-rich genomic regions during replication (39). With c-Myc activation being a crucial event in many human cancers, identifying the mechanisms through which this oncogene promotes RS provides critical insights into cancer biology and therapy.

## Methods

Sequencing data have been deposited in GEO under accession code GSE146766

### Cell culture and treatments

Cell lines used were immortalized hTERT human Retinal Pigment Epithelia 1 (ATCC CRL-4000) c-Myc-ER cells (previously described in (26)) and Retinal Pigment Epithelia 1 (ATCC CRL-4000) ER empty. Cells were cultured in phenol red free DMEM/F12 media supplemented with 10% charcoal-treated fetal bovine serum, 1% penicillin/streptomycin (Gibco) and 3% sodium bicarbonate (Gibco). Cells were maintained in Puromycin (2 *μ*g/ml). Cells were treated with 4-hydroxytamoxyfen (4OH-T) (100 nM), with HU overnight (2 mM) and for 4 hr (0.2 mM), with Nocodazole for 8 hr (200 ng/ml) and with Palbociclib for 24 hr (150 nM). RPE1 h-TERT (ATCC CRL-4000) cells were cultured in DMEM/F12 media supplemented with 10% fetal bovine serum, 1% penicillin/streptomycin (Gibco) and 3% sodium bicarbonate (Gibco). A549 (ATCC CCL-185) cells were cultured with DMEM/F12 media supplemented with 10% FBS and 1% penicillin/streptomycin (Gibco). H1299 (ATCC CRL-5803) cells were cultured in RPMI media supplemented with 10% FBS and 1% penicillin/streptomycin (Gibco).

### siRNA transfection

For siRNA transfection, Lipofectamine RNAiMAX (Invitrogen, 13778-075) was used following the manufacturer’s instructions. Experiments were carried out 20 hr after retro-transfection for synchronised cells. Unsynchronised cells were split 24 hr after transfection and then used for experiments 24 hr later. siRNA oligonucleotides with the following sequences, were used: siRad21 (GACCAAGGUUCCAUAUUAU), siCTCF (GGAGCCUGCCUGCCGUAGAAAUUTT), siMau2 (CCUCAGAACUUAACAUCUG). Non-targeting siRNA, referred as to sicont, (D-001206-13-05 siGENOME Non-Targeting siRNA pool) was purchased from Dharmacon.

### Plasmid transfection

For transient plasmid transfection, Lipofectamine 2000 (Invitrogen, 11668019) was used following the manufacturer’s instructions. Plasmids used were pEGFP N1 and Scc4-Flag-EGFP (34).

### Immunofluorescence

When appropriate cells were preextracted with ice-cold 0.2% triton solution in PBS 1x for 1 min and fixed for 20 min in formaldehyde solution 4%. If the extraction protocol was not carried out, cells where permeabilised for 4 min in 0.2% triton solution in PBS 1x after fixation. Cells were blocked in blocking solution (1% BSA, 0.2% Tween in PBS) for 1 hr at room temperature. The incubation with the primary antibodies anti-RPA2 (Millipore RPA34-20 1:500), anti-Phospho-Histone H2A.X (*γ*H2AX) (Ser139) (Cell Signaling Technology *γ*H2AX 20E3 1:250), anti-Smc1 (Bethyl laboratories A300-055A 1:1000), anti-Smc3 (Bethyl laboratories A300-060A 1:2000), anti-Mcm7 (Santa Cruz Biotechnology sc-56324 1:150), anti-RPA32 Phospho S4/S8 (Bethyl laboratories A300-245A 1:500) and anti-GFP (Abcam AB1218 1:1000) was carried out overnight. The following day the coverslips were incubated with secondary antibodies (Alexa Fluor^®^ 488 goat anti-mouse 1:2000 and Alexa Fluor^®^ 647 goat anti-rabbit 1:2000) for 2 hr at room temperature. The cells were stained with Hoechst (Invitrogen) solution 1:10000 in PBS 1x. The coverslips were mounted on slides with mounting medium Fluoroshied (Sigma). Images were obtained with Leica SPE2 40x objective lens and processed with Fiji. For the quantitative analysis, between 200 and 300 cells were analysed per sample.

### EdU and EU incorporation

Cells on coverslips were incubated with EdU (final concentration 10 μM) for 30 min and fixed in 4% formaldehyde solution. Cells were permeabilised in 0.2% triton for 5 min and incubated with Click-it reaction cocktail (Click-it Alexa Fluor 647 C-10424 Invitrogen) for 30 min. Nuclei were stained with Hoechst (Invitrogen) solution 1:10000. The coverslips were mounted on slides with mounting medium Fluoroshied (Sigma). Images were obtained with Leica SPE2 40x objective lens and processed with Fiji.

For detection of global RNA synthesis levels by 5-Ethynyl Uridine (EU) staining, 1 mM EU was added to cells for 1hr prior to collection. Cells were fixed in 4% formaldehyde. EU detection was performed using the Click-iT RNA Alexa Fluor 488 Imaging kit (Ther-moFisher, C10329) following manufacturer’s instructions. Coverslips were rinsed for 2 min in Click-iT reaction rinse buffer and stained in Hoechst solution (1:10,000, Invitrogen H3570) for 5 min at room temperature. Fluoroshield (Sigma, F6182) was used for mounting on slides. Once dry, coverslips were sealed with nail varnish. Leica SPE2 using 40x objective lens and processed with Fiji.

### Fibre analysis

Cells were labelled with 25μM CIdU for 15 min at 37°C and then with 250μM CO2-equilibrated IdU (final concentration 250 μM) for 15 min at 37°C. Fibre spreading and labelling was performed as in (40). The fibres were stained with primary antibodies (Rat anti-BrdU Abcam ab6326 1:250, Mouse anti-BrdU BD Biosciences 347580 1:100) overnight and with secondary antibodies (Alexafluor 555 goat anti-rat 1:500, Alexafluor 488 goat anti-mouse1:500) for 1.5 hr. Images were obtained with Leica SPE2 63x objective lens and processed with Fiji. 100-200 fibres were measured for each experiment. Composite images were constructed to visualise red and green channels simultaneously. The ‘line’ tool in Fiji was used to measure length of DNA replicating fibres, which are characterised by the presence of consecutive red and green signals. Total amount of fibres (ongoing fibres, replication origins, replication terminations, stalled forks) and replication origins (characterised by a red track between two green tracks) were counted to quantify the percentage of origin firing.

### Chromatin preparation

RPE-1 c-Myc ER cells were seeded in 10 cm dishes and c-Myc expression was activated upon tamoxifen treatment. Cells were harvested after 16 hr of c-Myc activation and the chromatin was isolated with Chromatin Extraction Kit (ab117152, Abcam) according to the manufacture’s protocol. The sonication was performed in a Diagenode Bioruptor^®^ sonicator using the program 10 min: 30 s on, 30 s off.

### Western blot

Cell extracts were prepared in RIPA buffer (Tris pH 7.5 20 mM, NaCl 150 mM, EDTA 1 mM, EGTA 1 mM, NP-40 1%, NaDoc 1%), phosphatase inhibitor cocktail 2 and 3 (Sigma P5726 and P0044) 1:1000, and protease inhibitor cocktail (Sigma P8340) 1:1000. Primary antibodies used anti-Phospho-Histone H2A.X (γH2AX) (Ser139) (Cell Signaling Technology g-H2AX 20E3 rabbit 1:250), anti-Scc4 (Mau2) (Abcam ab183033 1:500), anti-Idn3 (Nipbl) (Abcam ab106768 1:500), anti-Smc1 (Bethyl laboratories A300-055A rabbit 1:10000), anti-Smc3 (Bethyl laboratories A300-060A rabbit 1:10000), anti-Rad21 (Abcam ab992 1:2000), anti-CyclinE (Santa Cruz Biotechnology HE12 sc-247 1:1000), anti-CTCF (Chip-grade AB70303 Abcam 1:5000), anti-RPA32 Phospho S4/S8 (Bethyl laboratories A300-245A 1:1000), anti-RPA2 (Millipore RPA34-20 1:1000), anti-c-Myc (Santa Cruz Biotechnology 9E10 sc-40 1:1000), anti-CHK1 (Cell Signalling Technology 2360 1:1000), anti CHK1 Phospho S345 (Cell Signalling Technology 2341 1:250), anti-CHK2 (Cell Signalling Technology 2662 1:1000), anti CHK2 Phospho Thr68 (Cell Signalling Technology 2661 1:1000), anti-p21 (Cell Signalling Technology 2947 1:1000), anti-MDM2 (Cell Signalling Technology 86934 1:1000). Secondary antibodies used were goat anti-mouse IgG HRP conjugate (Thermofisher Scientific PA1-74421 1:4000) and goat anti rabbit IgG HRP conjugate (ThermoFisher Scientific 31460 1:4000). GAPDH (Genetex GT239 1:1000) and Vinculin (Abcam AB129002 1:10000) were used as loading controls.

### Flow cytometry

For analysis of DNA content by propidium iodide (PI) staining, cells were collected by trypsinisation and fixed in 70% ethanol at −20°C overnight. After centrifugation, the cell pellet was washed with PBS and resuspended in 100 mg/ml RNaseA and 50 mg/ml propidium iodide in PBS, and incubated at 4°C overnight. Samples were measured on a BD LSRII flow cytometer using DIVA software (BD) and analysed using FlowJo software.

### Survival assay

Cells were treated for 48 hr with 4OH-T or left untreated. The same volume of cell suspension was replated in 5 cm dishes and colonies were left to grow for one week. Cells were finally fixed and stained in 70% EtOH and 0.5% Methylene blue.

### CHiPseq

Cells were cultured in 15 cm dishes for 48 hr with or without the addition of 4OH-T. Cells were then washed with 1x PBS and crosslinked in 10ml of 1% formaldehyde at RT for 10 min. Quenching was carried out adding 1 ml of 1.25 M glycine for 10 min at RT. Cells were then scraped in PBS, spin down, resuspended in cold buffer A (100 mM Hepes pH8, 100 mM EDTA, 5 mM EGTA, 2.5% triton) and rocked for 10 min at 4°C. The same step was repeated using cold buffer B (100mM Hepes pH8,2M NaCl, 100mM EDTA, 5 mM EGTA, 0.1% triton). Cells were resuspended in cold ChIP buffer (25 mM tris/HCl pH8, 2 mM EDTA, 150 mM NaCl, 1% triton, 0.1% SDS) plus protease inhibitor cocktail (Sigma P8340) and sonicated at maximum output on a Bioruptor for 30 s on/30 s off for 30 min using Diagenode tubes. Sonication was checked on 1% agarose gel. After sonication, lysates where centrifuged for 15 min at maximum speed at 4°C. Protein A solution was prepared by resuspending beads in ChIP buffer (about 50%) 1 μg/μl BSA and rocking at 4°C for 15 min. The supernatant (soluble chromatin) was transferred in new tubes and pre-cleared adding blocked protein A solution and rocking for 2 hr at 4°C. Cleared soluble chromatin was centrifuged for 4 min at 4000rpm at 4°C. The supernatant was transferred in a new tube and 10 μl was saved as input. The soluble chromatin was incubated overnight with 10 μg of anti Smc1 (Bethyl laboratories A300-055A rabbit). The following day 20 μl protein A beads prepared as above, were added to chromatin, which was then rocked at 4°C for 2 hr. Beads were spin down for 2 min at 2000 rpm and washed with ChIP buffer, wash solution 1 (25 mM Tris/HCl pH8, 2 mM EDTA, 500 mM NaCl, 1% triton, 0.1% SDS), Wash solution 2 (250 mM LiCl, 1% NP40, 1% NaDOC, 1 mM EDTA, 10 mM Tris/Hcl pH 8) and twice with TE. TE was then removed and elution buffer (1% SDS, 100 mM NaHCO3) was added. All samples were incubated at 65°C overnight to reverse crosslinking. The day after, samples were purified using QIAquick PCR purification kit (QUIAGEN). The DNA was then diluted in ddH2O. QC was performed using the ThermoFisher Qubit and either the Agillent BioAnalyser or TapeStation. The DNA samples were normalised and prepared into Illumina compatible libraries using the KAPA HyperPrep kit according to the manufacturer’s instructions. The libraries were pooled to 4 nM and sequencing was performed on the HiSeq 4000 with at least 75 bp reads.

### RTqPCR

RNA was extracted using RNeasy plus mini kit Qiagen. Before column purification cell pellets were vortexed for 30 s in RLT buffer +1% β-mercaptoethanol. RT-qPCR was carried out using Mesa Blue mastermix (Eurogentec). All reactions were normalised to Gapdh as a control.

### Statistics

Statistical significance was analysed using Mann-Whitney test and Student’s t-test. When appropriate, S phase cells were defined as the portion of cells where RPA2 > 40 a.u. (Figure 1f, 1g), RPA2 > 50 a.u. (Figure 2a, S3a), RPA2 > 25 a.u. (Figure S1q, S1r) RPA2 > 18 a.u. (Figure S3b).

## ACKNOWLEDGEMENTS

We are grateful to Dr. F Uhlmann for advice. We thank the Crick Institute Advanced Sequencing Facility for the ChIPseq. We thank Dr. J-M. Peters for kindly providing GFP-Mau2 plasmid. We would also like to thank to Dr. F. Peri for help with statistical analysis. This work was supported by core funding to the MRC-UCL University Unit (Ref. MCEXG0800785) and funded by R.d.B.’s Cancer Research UK Programme Foundation Award.

---

**Fig. S1.**
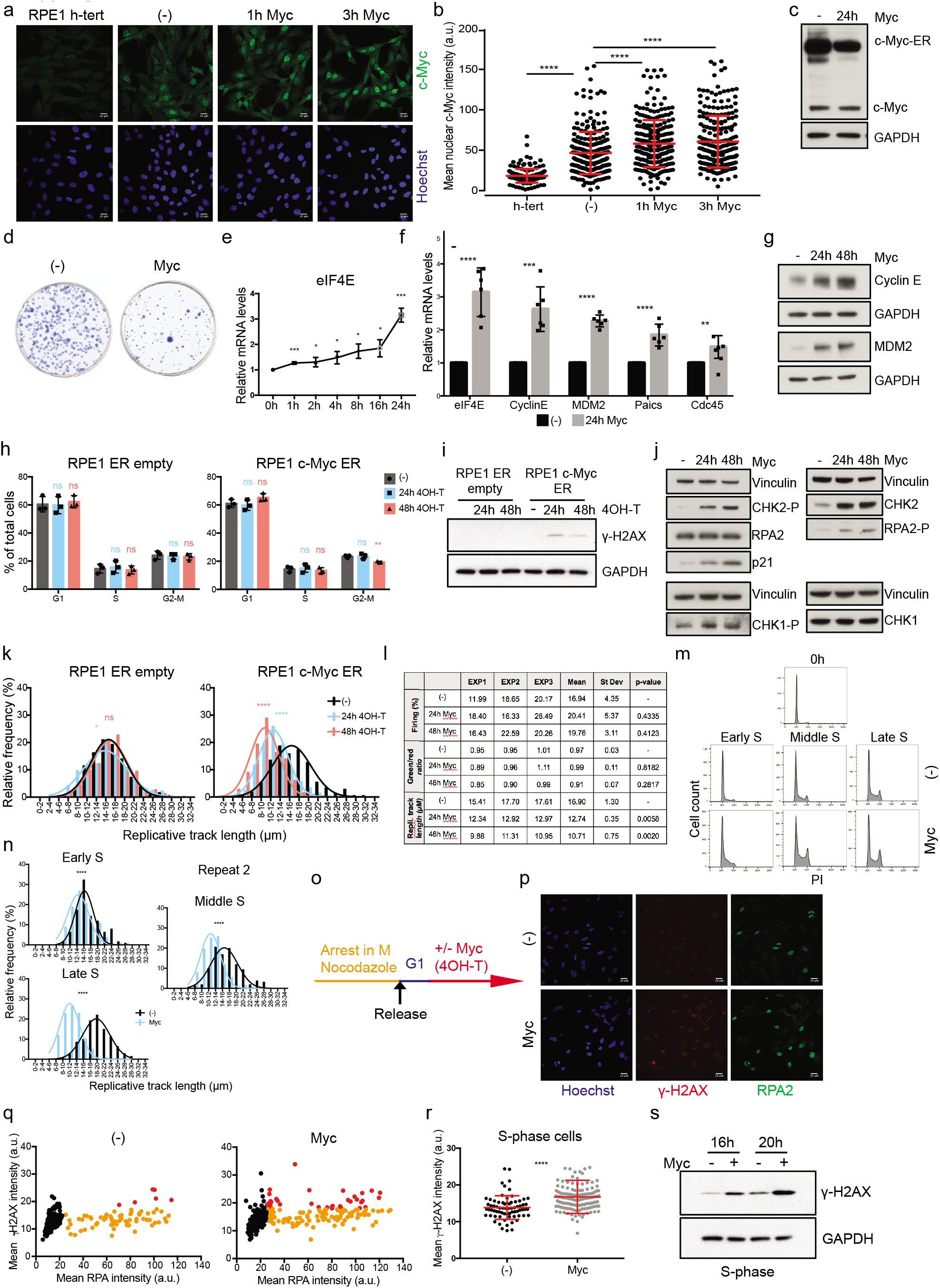
c-Myc-induced replication stress and DNA damage require G1 phase of the cell cycle. a) Immunofluorescence staining of c-Myc in RPE1 h-TERT and RPE1 c-Myc-ER cells after addition of 4OH-T for the indicated times. b) Graph showing c-Myc mean intensity in individual RPE1 h-TERT and RPE1 c-Myc ER cell nuclei after addition of 4OH-T for the indicated times, plotted in the scatter plot. p-value****<0.0001 calculated with Mann-Whitney test. n=1 experiment. c) Western blot of c-Myc in untreated cells and after 24 hr 4OH-T addition in RPE1 c-Myc ER cells. GAPDH is a loading control. Representative of n=3 experiments. d) Survival assay of untreated and c-Myc activated RPE1 c-Myc ER cells. Cells were treated with 48 hr with 4OH-T, plated in fresh media and colonies were left to grow for 7 days. Representative of n=4 experiments. e) Time course of the c-Myc target eIF4E mRNA levels at the indicated time-points after c-Myc activation. p-value***=0.0009 and 0.0006, *=0.02, 0.01, 0.01 calculated with Student’s t-test. n=4 experiments. f) mRNA levels of different c-Myc targets at 24 hr of c-Myc activation. p-value****<0.0001, ***=0.0002, **=0.0062 calculated with Student’s t-test. n=4 experiments. g) Western blot of the indicated c-Myc targets at 24 hr and 48 hr of c-Myc activation. GAPDH is a loading control. Representative of n=3 experiments. h) Cell cycle distribution determined by PI staining and FACS analysis of c-Myc ER and RPE1 ER empty cells with and without 4OH-T addition at the indicated time points. RPE1 c-Myc ER: p-value**=0.0038 calculated with Student’s t-test. n=3 experiments. i) Western blot of *γ*H2AX at the indicated time-points upon 4OH-T addition in RPE1 c-Myc ER and RPE1 ER empty cell lines. GAPDH is a loading control. Representative of n=3 experiments. j) Western blot of the indicated proteins at 24 hr and 48 hr of c-Myc activation. Vinculin is a loading control. Representative of n=3 experiments. k) Histograms reporting the distribution of fibre length for control and c-Myc-induced cells at the indicated times of c-Myc activation in RPE1 c-Myc ER and RPE1 ER empty cell lines; RPE1 c-Myc ER: p-value****<0.0001 calculated with Mann-Whitney test. RPE1 ER empty: p-value*=0.01 calculated with Mann-Whitney test. Pool of n=3 experiments. l) Table showing the percentage of origin firing, replication fork asymmetry (green/red ratio) and DNA fibre length for the experiments analysed in Figure S1k at 24 hr and 48 hr of c-Myc activation. Reported p-values were calculated with Student’s t-test. m) Cell cycle profile at the indicated time-points after release from G1 with and without c-Myc activation. Early: between 16 hr and 18 hr; middle: between 20 hr and 22 hr; late: 24 hr. n) Repeat of Figure 1d. Histograms reporting the distribution of fibre length for control and c-Myc-induced cells at the reported times after release from arrest; early=18 hr, middle=22 hr, late=24 hr; p-value****<0.0001 calculated with Mann-Whitney test. Repeat 2 of n=2 experiments. o) Schematic of the synchronisation experiments for G1 release with nocodazole. RPE1 c-Myc ER cells were treated with nocodazole for 8 hr. After mitotic shake-off cells were plated in fresh media. After cells were released into G1, 4OH-T was added to induce c-Myc or left untreated as control. p) Representative images of RPA and *γ*H2AX immunofluorescence. q) Immunofluorescence staining of chromatin-bound RPA and *γ*H2AX after nocodazole arrest. Scatter plot showing the intensity of RPA and *γ*H2AX signal in single nuclei. Black=RPA negative cells, orange=RPA positive cells, red=RPA positive cells with higher *γ*H2AX signal. n=1 experiment. r) Graph showing *γ*H2AX intensity in individual S phase cells plotted in the scatter plot. p-value****<0.0001 calculated with Mann-Whitney test. n=1 experiment. s) Western blot of *γ*H2AX at the indicated time-points after release from nocodazole arrest, with and without c-Myc activation. GAPDH is a loading control. Representative of n=2 experiments.

**Fig. S2.**
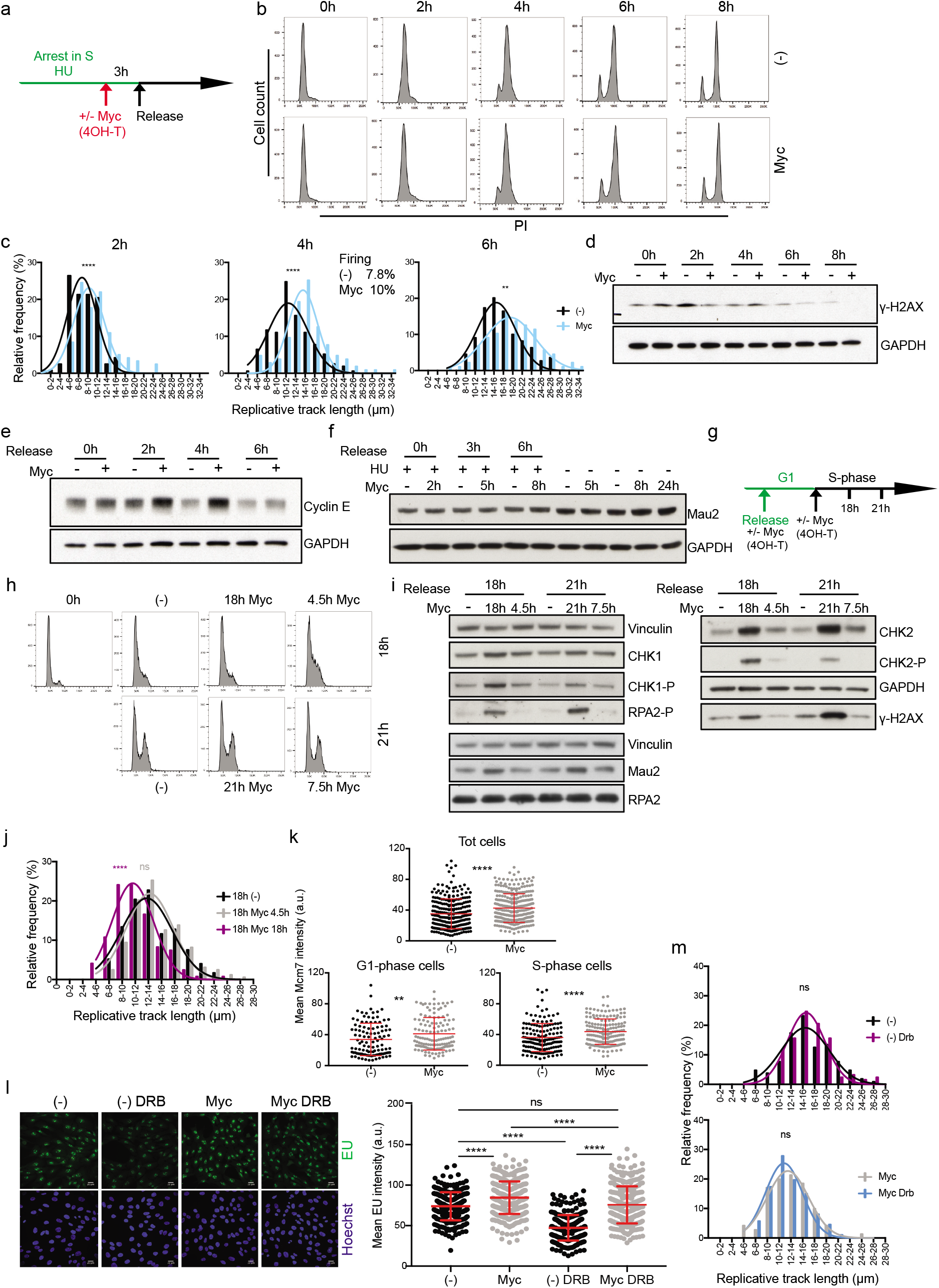
Replication initiation and replication-transcription collisions are not involved in generating replication stress and DNA damage upon c-Myc activation. a) Schematic of the synchronisation experiments for S release with HU. RPE1 c-Myc ER cells were arrested in HU overnight. 4-OHT was added 3 hr before release. Cells were then washed and released in S. b) Cell cycle profile at the indicated time-points after release from S with and without c-Myc activation. c) Histograms reporting the distribution of fibre length for control and c-Myc-induced cells at the reported times after release from HU arrest; p-value****<0.0001, **=0.006 calculated with Mann-Whitney test. n=1 experiment. d) Western blot of *γ*H2AX at the indicated time-points after release from HU-arrest, with and without c-Myc activation. GAPDH is a loading control. Representative of n=3 experiments. e) Western blot of Cyclin E at the indicated time-points after release from HU-arrest, with and without c-Myc activation. GAPDH is a loading control. Representative of n=3 experiments. f) Western blot of Mau2 at the indicated time-points after release from HU-arrest and in asynchrounous cells, with and without c-Myc activation for the indicated time-points. GAPDH is a loading control. Representative of n=3 experiments. g) Schematic of the synchronisation experiments for G1 release with Palbociclib. RPE1 c-Myc ER cells were arrested in G1 for 24 hr. Cells were then released and 4OH-T was added immediately upon release or 14 hr later, when cells enter S-phase. Cells were then collected at 18 hr and 21 hr after release. h) Cell cycle profile at the indicated time-points after release from Palbociclib G1-arrest with and without c-Myc activation for the indicated times. i) Western blot of the indicated proteins after release from Palbociclib G1-arrest, with and without c-Myc activation for the indicated times. GAPDH and Vinculin are loading controls. Representative of n=3 experiments. j) Histograms reporting the distribution of fibre length for control and c-Myc-induced cells at the reported times after release from Palbociclib G1-arrest; p-value****<0.0001 calculated with Mann-Whitney test. n=1 experiment. k) Graph showing chromatin-bound Mcm7 intensity in individual total, G1 and S phase pre-extracted nuclei after 18 hr from release from G1, plotted in the scatter plot. p-value****<0.0001 calculated with the Mann-Whitney test. n=1 experiment. l) Left: representative images of EU levels in control and c-Myc-induced cells with and without the addition of DRB for 2 hr. Right: graph showing EU intensity in individual cell nuclei plotted in the scatter plot at 20 hr after release from G1-arrest with and without the addition of DRB for 2 hr and c-Myc activation; p-value****<0.0001 calculated with Mann-Whitney test. Representative of n=3 experiments. m) Histograms reporting the distribution of fibre length for control and c-Myc-induced cells at 20 hr after release from G1-arrest with and without the addition of DRB for 2 hr; p-value calculated with Mann-Whitney test. Representative of n=3 experiments.

**Fig. S3.**
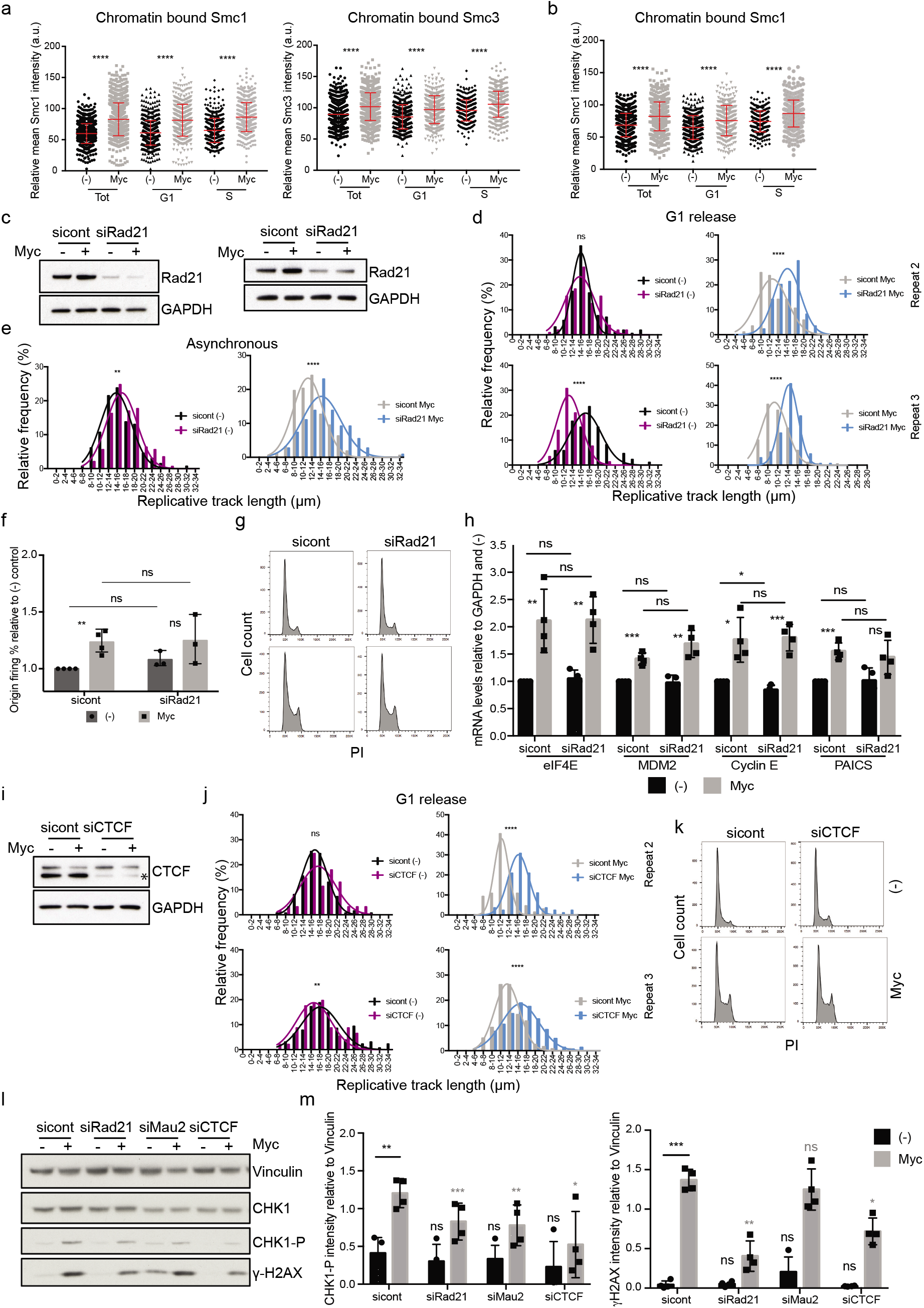
Reducing the levels of cohesin on chromatin or at CTCF sites prevents c-Myc induced replication stress without affecting the extent of origin firing and the cell cycle profiles. a) Synchronised cells were released into the cell cycle and immunofluorescence of chromatin-bound cohesin subunits Smc1 and Smc3 were performed at 14hr after release. Graphs reporting the intensity of Smc1 and Smc3 chromatin signals in total, S phase and G1 phase single nuclei of untreated and c-Myc cells. p-value ****<0.0001 calculated with Mann-Whitney test. Pool of n=3 experiments. b) Graphs reporting the intensity of Smc1 chromatin signals in total, S phase and G1 phase single nuclei in asynchronous population at 16 hr of c-Myc activation and in untreated cells; p-value****<0.0001 calculated with Mann-Whitney test. Pool of n=3 experiments. c) Western blot showing Rad21 knock down in synchronous (left) and asynchronous (right) cells. GAPDH is a loading control. d) Histograms reporting the distribution of fibre length in synchronised sicontrol and siRad21 depleted cells. p-value****<0.0001 calculated with Mann-Whitney test. Repeats 2 and 3, Repeat 1 is in Figure 3d. e) Histograms reporting the distribution of fibre length in asynchronous sicontrol and siRad21 depleted cells. p-value****<0.0001, **=0.0026 calculated with Mann-Whitney test. Representative of n=3 experiments. f) Percentages of origin firing at 20 hr after release from G1-arrest and Rad21 or sicontrol knock down, with and without c-Myc activation. p-value**=0.0035 calculated with Student’s t-test. n=3 experiments. g) Cell cycle profile at 20 hr after release from G1 and Rad21 or sicontrol knock down, with and without c-Myc activation. Representative of n=3 experiments. h) mRNA levels of different c-Myc targets in control and Rad21 depleted cells at 24 hr of c-Myc activation. p-value calculated with Student’s t-test. n=4 experiments. i) Western blot showing CTCF knock down in synchronous cells. GAPDH is a loading control. j) Histograms reporting the distribution of fibre length in synchronised sicontrol and siCTCF depleted cells. p-value****<0.0001, **=0.0078 calculated with Mann-Whitney test. Repeats 2 and 3, repeat 1 is in Figure 3h. k) Cell cycle profile at 20 hr after release from G1 and CTCF or sicontrol knock down, with and without c-Myc activation. Representative of n=3 experiments. l) Western blot of the indicated proteins in control, Rad21, CTCF and Mau2 depleted cells upon 24 hr of c-Myc activation. Vinculin is a loading control. Representative of n=4 experiments. m) Western blot quantification of the indicated proteins in control, Rad21, CTCF and Mau2 depleted cells upon 24 hr of c-Myc activation relative to Vinculin. n=4 experiments.

**Fig. S4.**
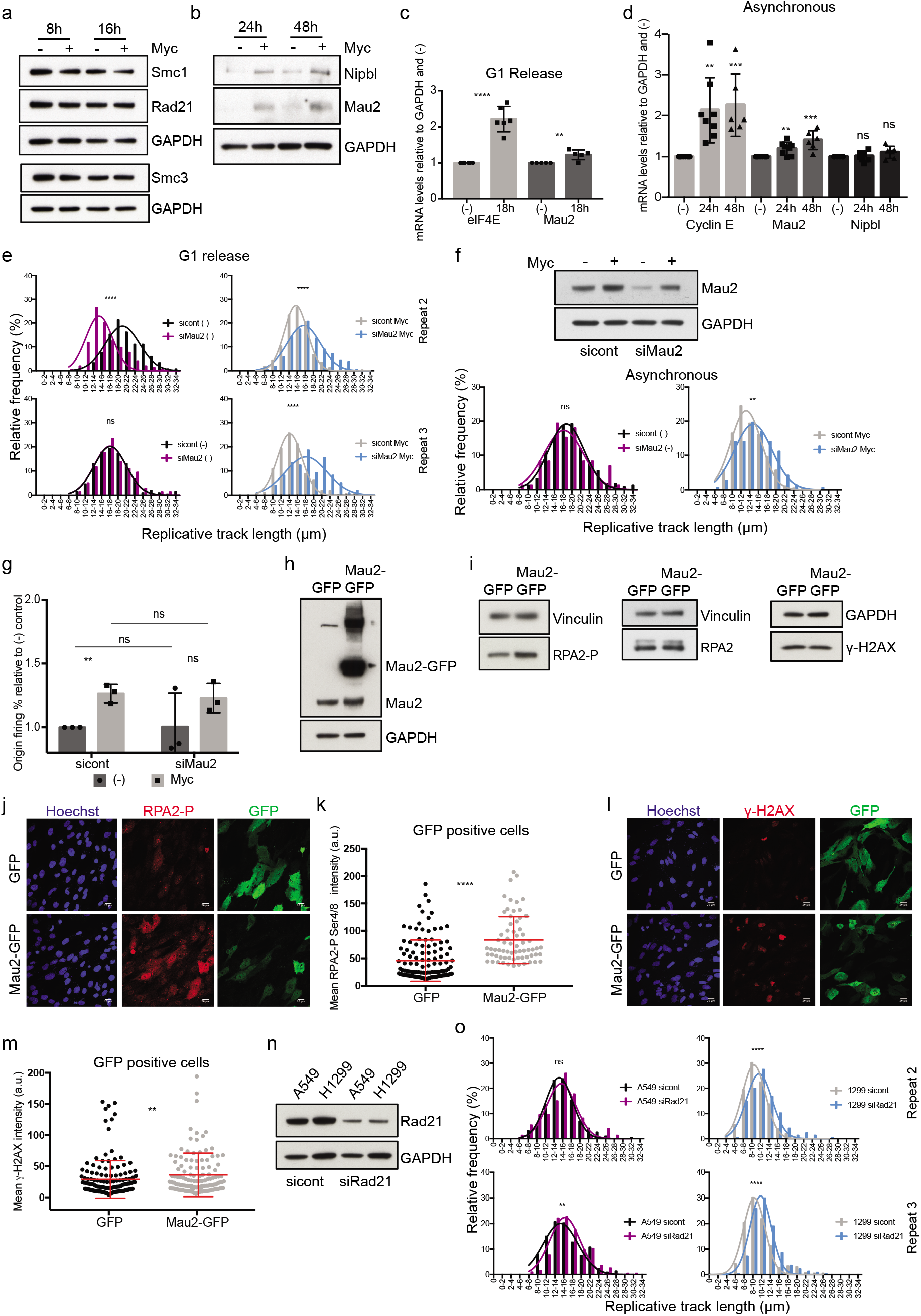
Dysregulation of the cohesin loader Mau2 is required for c-Myc-induced replication stress. a) Western blot showing total levels of cohesin subunits and regulators in asynchronous population with and without c-Myc activation for the indicated time-points. Representative experiment of n=2. b) Western blot showing total levels of the cohesin loaders Mau2 and Nipbl in asynchronous cells at 24 hr and 48 hr of c-Myc activation. Representative experiment of n=3. c) mRNA levels of Mau2 at 18 hr of c-Myc activation in synchronised cells. p-value****<0.0001, **=0.0056 calculated with Student’s t-test. n=6 experiments. d) mRNA levels of Mau2 and NIPBL at the indicated time-points of c-Myc activation in asynchronous population. p-value***= 0.0005, 0.0003, **= 0.0012, 0.0042 calculated with Student’s t-test. n=6 experiments. e) Histograms reporting the distribution of fibre length in synchronised sicontrol and siMau2 depleted cells. p-value****<0.0001 calculated with Mann-Whitney test. Repeats 2 and 3, repeat 1 is in 4g. f) Top: Western blot showing Mau2 knock-down in asynchronous population. GAPDH is a loading control. Bottom: histograms reporting the distribution of fibre length in asynchronous sicontrol and siMau2 depleted cells. p-value**=0.0057 calculated with Mann-Whitney test. n=1 experiment. g) Percentages of origin firing at 20 hr after release from G1-arrest and Mau2 knock-down, with and without c-Myc activation. p-value**=0.0034 calculated with Student’s t-test. n=3 experiments. h) Western blot showing total levels of Mau2 in RPE1 h-TERT cells after transient transfection with GFP and Mau2-GFP plasmids for 24 hr. Representative of n=4 experiments. i) Western blot showing total levels of the indicated proteins in RPE1 h-TERT cells after transient transfection with GFP and Mau2-GFP plasmids for 24 hr. Vinculin and GAPDH are loading controls. Representative of n=4 experiments. j) Representative images of GFP and RPA2 phospho S4/8 levels after transient transfection with GFP and Mau2-GFP plasmids for 24 hr. k) Graph showing RPA2 phospho S4/8 intensity in individual cell nuclei after transient transfection with GFP and Mau2-GFP plasmids for 24 hr plotted in the scatter plot. Only GFP positive cells are shown. p-value****<0.0001 calculated with Mann-Whitney test. Representative of n=3 experiments. l) Representative images of GFP and *γ*H2AX levels after transient transfection with GFP and Mau2-GFP plasmids for 24 hr. m) Graph showing *γ*H2AX intensity in individual cell nuclei after transient transfection with GFP and Mau2-GFP plasmids for 24 hr plotted in the scatter plot. Only GFP positive cells are shown. p-value**=0.0074 calculated with Mann-Whitney test. Representative of n=3 experiments. n) Western blot showing Rad21 knock down in A549 and H1299 cells. GAPDH is a loading control. o) Histograms reporting the distribution of fibre length in sicontrol and siRad21 depleted A549 and H1299 cells. p-value****<0.0001, **=0.0031 calculated with Mann-Whitney test. Repeats 2 and 3, repeat 1 is in 4k.

## Notes

### Competing Interest Statement

The authors have declared no competing interest.

